# Late Pleistocene human genome suggests a local origin for the first farmers of central Anatolia

**DOI:** 10.1101/422295

**Authors:** Michal Feldman, Eva Fernández-Domínguez, Luke Reynolds, Douglas Baird, Jessica Pearson, Israel Hershkovitz, Hila May, Nigel Goring-Morris, Marion Benz, Julia Gresky, Raffaela A. Bianco, Andrew Fairbairn, Gökhan Mustafaoğlu, Philipp W. Stockhammer, Cosimo Posth, Wolfgang Haak, Choongwon Jeong, Johannes Krause

**Affiliations:** Department of Archaeogenetics, Max Planck Institute for the Science of Human History (MPI-SHH), Kahlaische Strasse 10, D-07745, Jena, Germany.; Department of Archaeology, Durham University, Durham, South Road, DH1 3LE, United Kingdom.; School of Natural Sciences and Psychology, Liverpool John Moores University, Liverpool, Byrom Street, L3 3AF, United Kingdom.; Department of Archaeology, Classics and Egyptology, 8-14 Abercromby Square, University of Liverpool, L69 7WZ, United Kingdom.; Department of Anatomy and Anthropology, The Dan David Center for Human Evolution and Biohistory Research and The Shmunis Family Anthropology Institute, Sackler Faculty of Medicine, Tel Aviv University, Post Office Box 39040, Tel Aviv 6997801, Israel.; The Steinhardt Museum of Natural History, Tel Aviv University, Post Office Box 39040, Tel Aviv 6997801, Israel; Department of Prehistory, Institute of Archaeology, The Hebrew University of Jerusalem, Jerusalem 91955, Israel.; Department of Near Eastern Archaeology, Free University Berlin, Fabeckstrasse 23-25, 14195 Berlin.; Department of Natural Sciences, German Archaeological Institute, 14195 Berlin, Germany.; School of Social Science, The University of Queensland, Michie Building, St Lucia, Brisbane, Queensland, Australia.; Department of Archaeology, Zonguldak Bülent Ecevit University, 67100 Incivez, Zonguldak, Turkey.; Institut für Vor- und Frühgeschichtliche Archäologie und Provinzialrömische Archäologie Ludwig-Maximilians-Universität München München, Schellingstrasse 12, D-80799, Germany.

## Abstract

Anatolia was home to some of the earliest farming communities. It has been long debated whether a migration of farming groups introduced agriculture to central Anatolia. Here, we report the first genome-wide data from a 15,000-year-old Anatolian hunter-gatherer and from seven Anatolian and Levantine early farmers. We find high genetic continuity (∼80-90%) between the hunter-gatherer and early farmers of Anatolia and detect two distinct incoming ancestries: an early Iranian/Caucasus related one and a later one linked to the ancient Levant. Finally, we observe a genetic link between southern Europe and the Near East predating 15,000 years ago that extends to central Europe during the post-last-glacial maximum period. Our results suggest a limited role of human migration in the emergence of agriculture in central Anatolia.

---

The practice of agriculture began in the Fertile Crescent of Southwest Asia as early as 10,000 to 9,000 BCE. Subsequently, it spread across western Eurasia while increasingly replacing local hunting and gathering subsistence practices, reaching central Anatolia by c. 8300 BCE^*1-3*^.

Recent genetic studies have shown that in mainland Europe, farming was introduced by an expansion of early farmers from Anatolia that replaced much of the local populations^*4,* *5*^. Such mode of spread is often referred to as the demic diffusion model. In contrast, in regions of the Fertile Crescent such as the southern Levant and the Zagros Mountains (located between present-day eastern Iraq and western Iran) the population structure persists throughout the Neolithic transition^*6*^, indicating that the hunter-gatherers of these regions locally transitioned to a food producing subsistence strategy.

Central Anatolia has some of the earliest evidence of agricultural societies outside the Fertile Crescent^3^ and thus is a key region in understanding the early spread of farming. While archaeological evidence points to cultural continuity in central Anatolia^*3*^, due to the lack of genetic data from pre-farming individuals it remains an open question whether and to what scale the development of the Anatolian Neolithic involved immigrants from earlier farming centers admixing with the local hunter-gatherers.

Here, we report new genome wide data from eight prehistoric humans (Fig.1A, Table 1, table S1), including the first Epipalaeolithic Anatolian hunter-gatherer sequenced to date (labeled AHG; directly dated to 13,642-13,073 BCE, excavated from the site of Pinarbaşi, Turkey), 5 early Neolithic Aceramic Anatolian farmers (labeled AAF; c. 8300-7800 cal BCE, one directly dated to 8269-8210 cal BCE^*3*^, from the site of Boncuklu, Turkey), adding to previously published genomes from this site^*7*^, and two Early Neolithic (PPNB) farmers from the southern Levant (One labeled KFH2, directly dated to c. 7,700-7,600 BCE; from the site of Kfar HaHoresh, Israel and the second labeled BAJ001, c. 7027-6685 BCE, from the site of Ba’ja, Jordan). This data comprises a genetic record stretching from the Epiplaeolithic into the Early Holocene, spanning the advent of agriculture in the region.

**Table 1.** An overview of ancient genomes reported in this study. For each individual the analysis group is given (AHG = Anatolian hunter-gatherer; AAF = Anatolian Aceramic early farmer; Levant_Neol = Levantine early farmer). When ^14^C dating results are available the date is given in cal BCE in 2-sigma range, otherwise a date based on the archaeological context is provided (detailed dating information is provided in Supplementary text S1 and table S1). The proportion of human DNA and the mean coverage on 1240K target sites in the ‘1240K’ enriched libraries are given. Uniparental haplogroups (mt = mitochondrial; Ychr = Y chromosome) are listed. Detailed information on the uniparental analysis can be found in Supplementary text S1 and data table S6.

| ID | Library name | Analysis group | Estimated date | Site | Sampled tissue | Total sequenced reads ($\times 10^6$) | Human DNA (%) | Mean Coverage (fold) | Genetic sex | mt | Ychr |
| --- | --- | --- | --- | --- | --- | --- | --- | --- | --- | --- | --- |
| ZBC | IPB001.B/C0101 | AHG | 13,642-13,073 cal BCE | Pınarbaşı | Intermediate phalanx | 126.7 | 33 | 2.9 | Male | K2b | C1a2 |
| ZHAG | BON004.A0101 | AAF | 8300-7800 BCE | Boncuklu | Petrous | 92.0 | 38 | 1.48 | Female | N1a1a1 |  |
| ZMOJ | BON014.A0101 | AAF | 8300-7800 BCE | Boncuklu | Third molar | 77.9 | 27 | 0.8 | Male | K1a | C |
| ZKO | BON001.A0101 | AAF | 8300-7800 BCE | Boncuklu | Petrous | 84.8 | 31 | 0.9 | Male | U3 | G2a2b2b |
| ZHJ | BON024.A0101 | AAF | 8300-7800 BCE | Boncuklu | Third molar | 87.7 | 38 | 0.76 | Female | U3 |  |
| ZHAJ | BON034.A0101 | AAF | 8269-8210 cal BCE | Boncuklu | Petrous | 75.4 | 30 | 0.69 | Female | U3 |  |
| KFH2 | KFH002.A0101 | Levant_Neol | 7712-7589 cal BCE | Kfar HaHoresh | Petrous | 342.0 | 8 | 0.16 | Female | N1a1b |  |
| BAJ001 | BAJ001.A0101 | Levant_Neol | 7027-6685 cal BCE | Ba'ja | Petrous | 17.3 | 45 | 0.75 | Female | N1b1a |  |

By analyzing this data, we find that the Anatolian hunter-gatherers are genetically distinct from other reported late Pleistocene populations and thus represent a previously undescribed population. We reveal that Neolithic Anatolian populations derive a large fraction of their ancestry from the Epipaleolithic Anatolian population, suggesting farming was adopted locally by the hunter-gatherers of central Anatolia. We also detect distinct genetic interactions between the populations of central Anatolia and earlier farming centers to the east, during the late Pleistocene/early Holocene as well as with European hunter-gatherers to the west during the Late Pleistocene.

## Results

We extracted DNA from the ancient human remains and prepared it for next-generation sequencing^*8,* *9*^ which resulted in human DNA yields lower than 2% (data table S1), comparable with low DNA preservation previously reported in the region^*6,* *7*^. To generate genome wide data, despite the low DNA yields we performed in-solution DNA enrichment targeting 1.24 million genome-wide single nucleotide polymorphisms (SNPs) (‘1240k capture’)^*10*^, which resulted in 129,406 to 917,473 covered SNPs per individual. We estimated low mitochondrial contamination levels for all eight individuals (1-6%; Materials and Methods and table S2) and could further test the males for nuclear contamination, resulting in low estimates (0.05-2.23%; table S2). For population genetic analyses, we merged genotype data of the new individuals with previously published datasets from 587 ancient individuals and 254 present-day populations (data table S2).

To estimate how the ancient individuals relate to the known west Eurasian genetic variation, we projected them onto the top two dimensions of present-day principal component analysis (PCA)^*6*^ (Fig. 1B). Strikingly, the AHG individual is positioned near both AAF and later Anatolian Ceramic farmers^*10*^ (labeled ACF; 7,000 - 6,000 cal BCE) with a slight leftward shift. These three prehistoric Anatolian populations (AHG, AAF and ACF), that represent a temporal transect spanning the transition into farming, are positioned between Mesolithic western European hunter-gatherers (WHG)^*4,* *10,* *11*^ at the far left and Levantine Epipalaeolithic Natufians^*6*^ at the far right. The newly reported Neolithic farmers^*6*^ (BAJ001 and KFH2) are positioned near the published ones^*6*^ (Supplementary Text S2). In ADMIXTURE analysis, AHG, AAF and ACF are all modeled as a mixture of two components that are each maximized in Natufians and WHG, consistent with their positions in PCA (fig. S1).

**Fig. 1.**
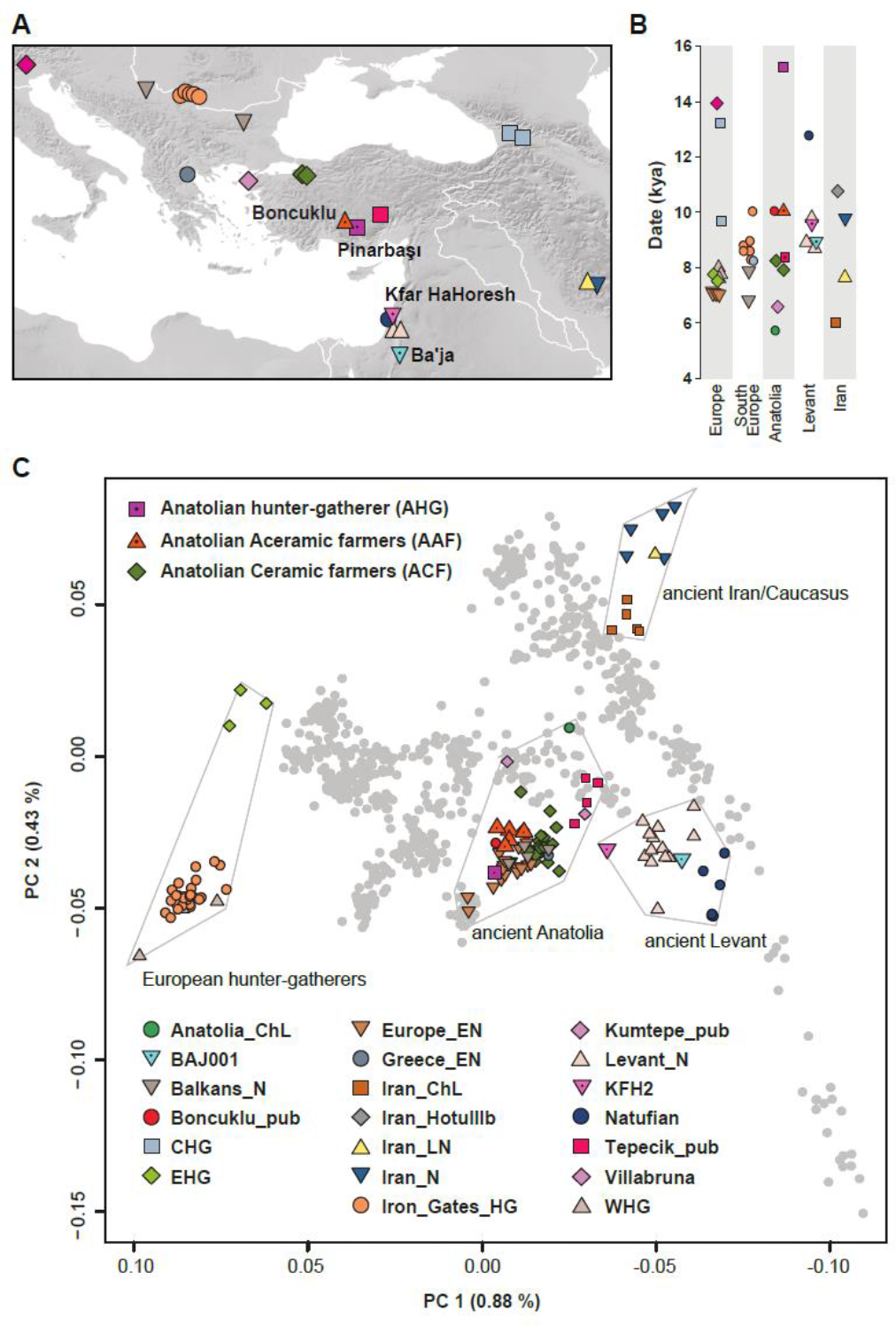
Location, age and PCA of analyzed individuals. **(A)** Locations of newly reported and selected published genomes. Archaeological sites from which new data is reported are annotated. Symbols for the analyzed groups are annotated in (C). (**B)** Average ages of ancient groups. (**C)** Ancient genomes (marked with color-filled symbols) projected onto the principal components computed from present-day west Eurasians (grey circles) (fig. S4). The geographic location of each ancient group is marked in (A). Ancient individuals newly reported in this study are additionally marked with a black dot inside the symbol.

Based on our observations in PCA and ADMIXTURE analysis we formally tested the ancestral compositions of the three Anatolian populations. We first characterized the ancestry of AHG. As expected from AHG’s intermediate position on PCA between Epipaleolithic/Neolithic Levantines and WHG, Patterson’s D-statistics^*12*^ of the form *D (AHG, WHG; Natufian/Levant_N, Mbuti)* ≥ 4.8 SE (standard error) and *D (AHG, Natufian/Levant_N; WHG, Mbuti)* ≥ 9.0 SE (table S3) indicates that AHG is distinct from both the WHG and Epipaleolithic/Neolithic Levantine populations and yet shares extra affinity with each when compared to the other. Accordingly, we find an adequate two-way admixture model using *qpAdm*^*12*^ (*χ*^*2*^*p* = 0.158), in which AHG derives around half of his ancestry from a Neolithic Levantine-related gene pool (48.0 ± 4.5 %; estimate ± 1 SE) and the rest from the WHG-related one (tables S4 and S5). These results support a late Pleistocene presence of both ancestries in a mixed form in central Anatolia. Notably, the genetic connection with the Levant predates the advent of farming in this region by at least five millennia and potentially correlates with evidence of human interactions between central Anatolia and the Levant during the Epipalaeolithic^*13*^.

In turn, AAF is slightly shifted upwards compared to AHG in the PCA, to the direction where ancient and modern Caucasus and Iranian groups are located. Likewise, when compared to AHG by *D(AAF, AHG; test, Mbuti)*, the AAF early farmers show extra affinity with early Holocene populations from Iran or Caucasus and with present-day South Asians, who have also been genetically linked with Iranian/Caucasus ancestry^*14,* *15*^ (Fig. 2A, fig. S2 and data table S3). A mixture of AHG and Neolithic Iranians provides a good fit to AAF in our *qpAdm* modeling (*χ*^*2*^*p* = 0.296), in which they derive most of their ancestry (89.7 ± 3.9 %) from a population related to AHG (tables S4 and S6). This suggests a long-term genetic stability in central Anatolia over five millennia despite changes in climate and subsistence strategy. Given that our admixture model for AHG does not require the Neolithic Iranian ancestry, it presumably diffused into central Anatolia during the final stages of the Pleistocene or early Holocene, most likely via contact through eastern Anatolia. This provides evidence of interactions between eastern and central Anatolia in the Younger Dryas or the first millennium of the Holocene, currently poorly documented archaeologically.

**Fig. 2.**
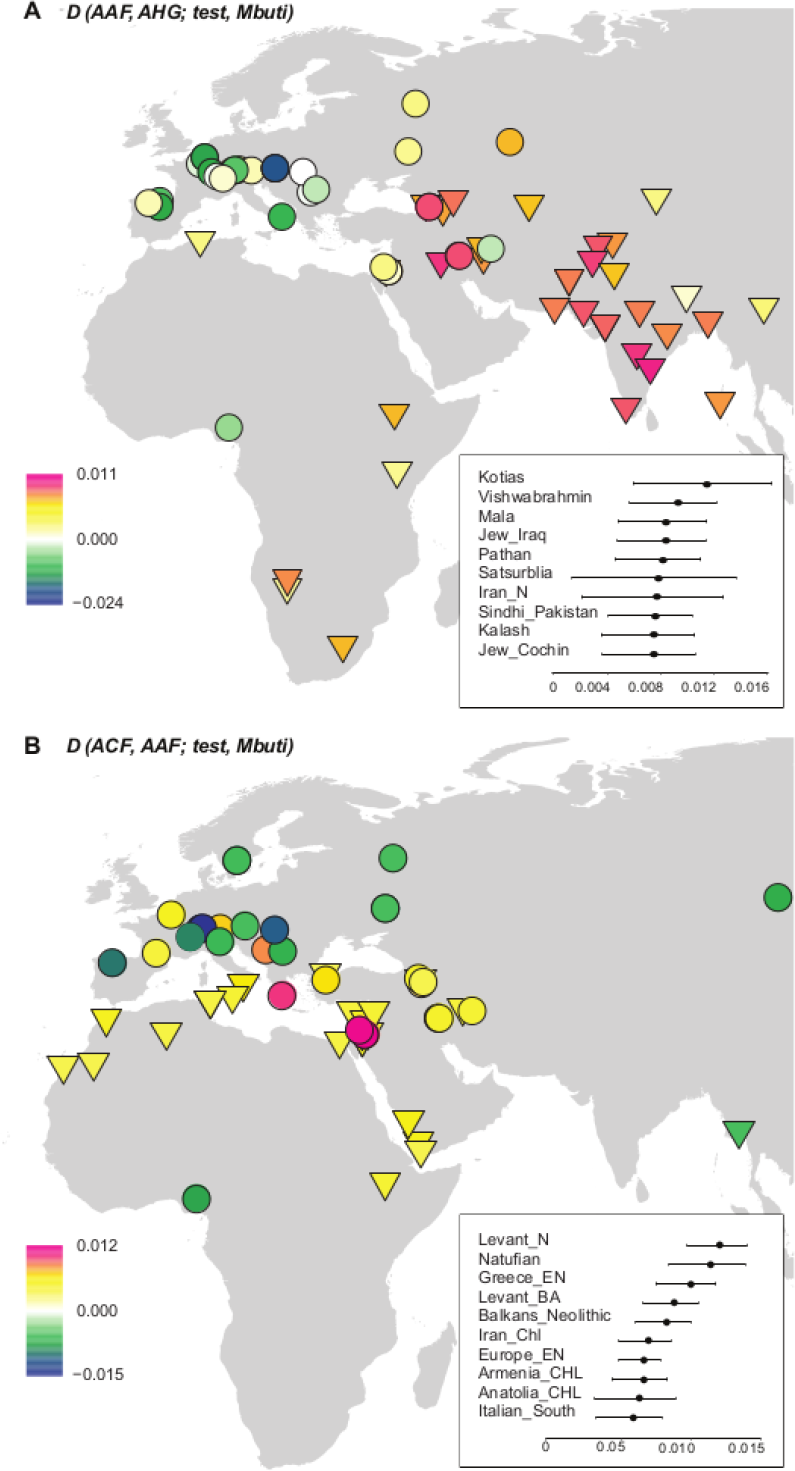
Differences between the ancient Anatolian populations in their genetic affinity to worldwide present-day and ancient populations. We plot the highest and lowest 40 values of *D(population 1, population 2; test, Mbuti)* on the map. Circles mark ancient populations and triangles present-day ones. “test” share more alleles with *population 1* when values are positive and with *population 2* when negative. The statistics and SEs are found in figs S2-S3 and data table S3. (**A)** Early Holocene Iranian and Caucasus populations, as well as present-day South Asians, share more alleles with AAF than AHG, measured by positive *D(AAF, AHG; test, mbuti)*. The top 10 values with ±1 SE are shown in the upper box. (**B)** Ancient Levantine populations share more alleles with ACF than AAF, measured by positive *D(ACF, AAG; test, Mbuti)*. The top 10 values with ±1 SE are shown in the lower box.

In contrast, we find that the later ACF individuals share more alleles with the early Holocene Levantines than AAF do, as shown by positive *D(ACF, AAF; Natufian/Levant_N, Mbuti)* ≥ 3.84 SE (Fig. 2B, fig. S3 and data table S3). Ancient Iran/Caucasus populations and contemporary South Asians do not share more alleles with ACF (|D| < 3.3 SE). Likewise, qpAdm modeling suggests that the AAF gene pool still constitutes more than 3/4 of the ancestry of ACF 2,000 years later (78.7 ± 3.5 %; tables S4 and S7) with additional ancestry well modeled by the Neolithic Levantines (*χ*^*2*^*p* = 0.115) but not by the Neolithic Iranians (*χ*^*2*^*p* = 0.076; the model estimated infeasible negative mixture proportions) (tables S4 and S7). These results suggest gene flow from the Levant to Anatolia during the early Neolithic. In turn, Levantine early farmers (Levant_Neol) that are temporally intermediate between AAF and ACF could be modeled as a two-way mixture of Natufians and AHG or AAF (18.2 ± 6.4 % AHG or 21.3 ± 6.3 % AAF ancestry; tables S4 and S8 and data table S4), confirming previous reports of an Anatolian-like ancestry contributing to the Levantine Neolithic gene pool^6^. These two distinct detected gene flows support a reciprocal genetic exchange between the Levant and Anatolia during the early stages of the transition to farming.

Anatolian hunter-gatherers experienced climatic changes during the last glaciation^*16*^ and inhabited a region that connects Europe to the Near East. However, interactions between Anatolia and Southeastern Europe in the later Upper Palaeolithic/Epipalaeolithic are so far not well documented archaeologically. Interestingly, a previous genomic study showed that present-day Near-Easterners share more alleles with European hunter-gatherers younger than 14,000 BP (‘Later European HG’) than with earlier ones (‘Earlier European HG’)^*17*^. With ancient genomic data available, we could directly compare the Near-Eastern hunter-gatherers (AHG and Natufian) with the European ones. As is the case for present-day Near-Easterners, the Near-Eastern hunter-gatherers share more alleles with the Later European HG than with the Earlier European HG, shown by the significantly positive statistic *D(Later European HG, Earlier European HG; AHG/Natufian, Mbuti*) (Fig. 3A and data table S5). Among the Later European HG, recently reported Mesolithic hunter-gatherers from the Balkan peninsula, which geographically connects Anatolia and central Europe (‘Iron Gates HG’)^*18*^, are genetically closer to AHG when compared to all the other European hunter-gatherers, as shown in the significantly positive statistic *D(Iron_Gates_HG, European hunter-gatherers; AHG, Mbuti/Altai)*. Iron Gates HG are followed by Epigravettian and Mesolithic individuals from Italy and France (Villabruna14 and Ranchot88 respectively^*17*^) as the next two European hunter-gatherers genetically closest to AHG (Fig. 3A and data table S5). Iron Gates HG have been suggested to be genetically intermediate between WHG and eastern European hunter-gatherers (EHG) with an additional unknown ancestral component^*18*^. We find that Iron Gates HG can be modeled as a three-way mixture of Near-Eastern hunter-gatherers (25.8 ± 5.0 % AHG or 11.1 ± 2.2 % Natufian), WHG (62.9 ± 7.4 % or 78.0 ± 4.6 % respectively) and EHG (11.3 ± 3.3 % or 10.9 ± 3 % respectively); (tables S4 and S9). The affinity detected by the above D-statistic can be explained by gene flow from Near-Eastern hunter-gatherers into the ancestors of Iron Gates or by a gene flow from a population ancestral to Iron Gates into the Near-Eastern hunter-gatherers as well as by a combination of both. To distinguish the direction of the gene flow, we examined the Basal Eurasian ancestry component (α), which is prevalent in the Near East ^*6*^ but undetectable in European hunter-gatherers^*17*^. Following a published approach^*6*^, we estimated α to be 24.8 ± 5.5 % in AHG and 38.5 ± 5.0 % in Natufians (Fig. 3B, table S10), consistent with previous estimates for the latter^*6*^. Under the model of unidirectional gene flow from Anatolia to Europe, 6.4 % is expected for α of Iron Gates by calculating (% AHG in Iron Gates HG) × (α in AHG). However, Iron Gates can be modeled without any Basal Eurasian ancestry or with a non-significant proportion of 1.6 ± 2.8 % (Fig. 3B, table S10), suggesting that unidirectional gene flow from the Near East to Europe alone is insufficient to explain the extra affinity between the Iron Gates HG and the Near-Eastern hunter-gatherers. Thus, it is plausible to assume that prior to 15,000 years ago there was either a bidirectional gene flow between populations ancestral to Southeastern Europeans of the early Holocene and Anatolians of the late glacial or a dispersal of Southeastern Europeans into the Near East. Presumably, this Southeastern European ancestral population later spread into central Europe during the post-last-glacial maximum (LGM) period, resulting in the observed late Pleistocene genetic affinity between the Near East and Europe.

**Fig. 3.**
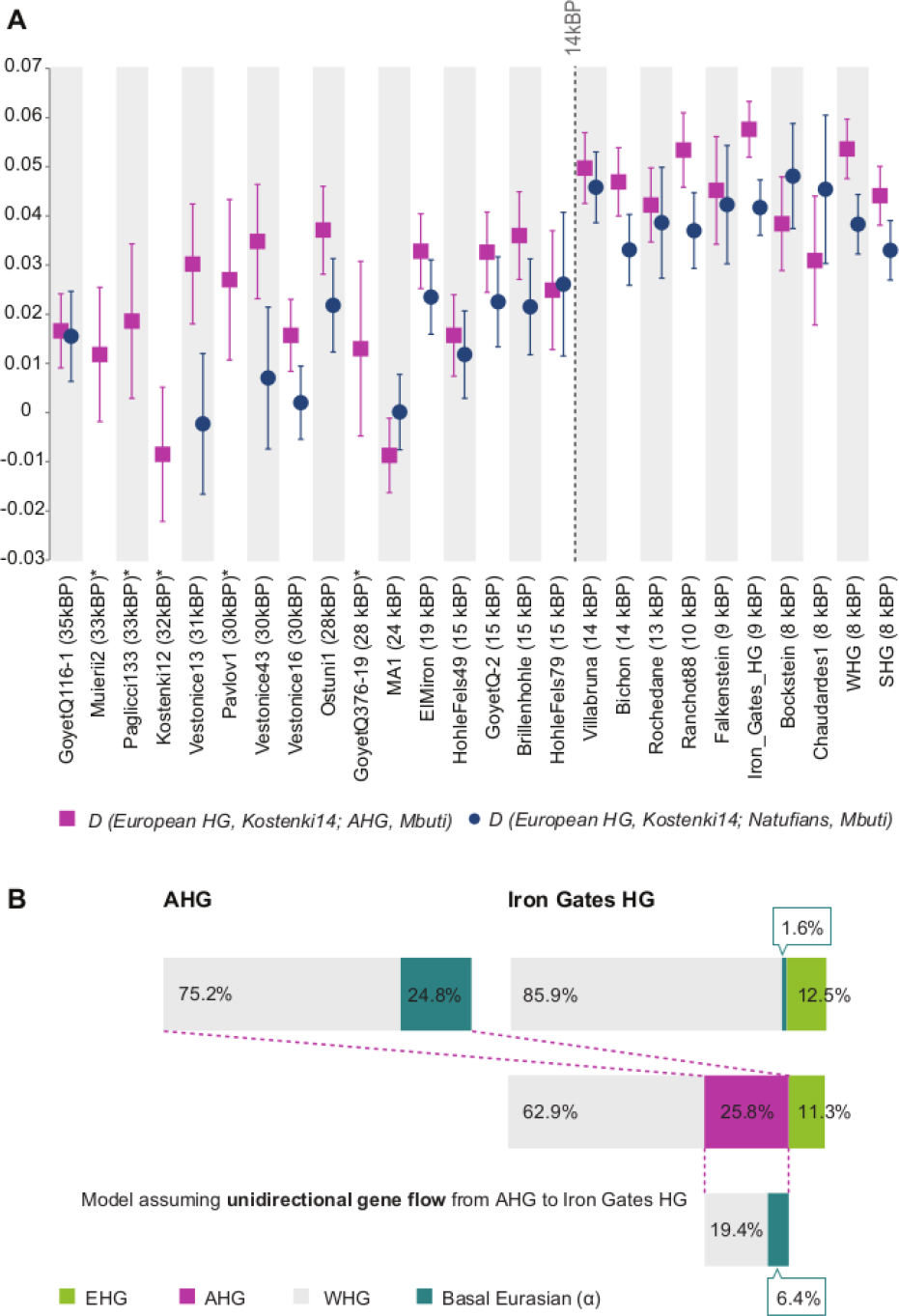
Genetic links between Near-Eastern and European hunter-gatherers. **(A)** Genetic affinity between Near-Eastern and European hunter-gatherers increases after 14,000 years ago as measured by the statistic *D(European HG, Kostenki14; Natufian/AHG, Mbuti)* Vertical lines mark ±1 SE. *Kostenki14* serves here as a baseline for the earlier European hunter-gatherers. Statistics including all analyzed European hunter-gatherers are listed in data table S5. Individuals marked with an asterisk did not reach the analysis threshold of over 30,000 SNPs overlapping with *Natufian/AHG*. (**B)** Basal Eurasian ancestry proportions (α) as a marker for Near-Eastern gene flow. Mixture proportions inferred by qpAdm for AHG and the Iron Gates HG are schematically represented ^*6*^. The lower schematic shows the expected α in Iron Gates HG under assumption of unidirectional gene flow, inferred from α in the AHG source population. The observed α for Iron Gates HG is considerably smaller than expected thus, the unidirectional gene flow from the Near East to Europe is not sufficient to explain the above affinity.

The uniparental marker analysis placed AHG within the mitochondrial sub-haplogroup K2b and within the Y-chromosome haplogroup C1a2, both rare in present-day Eurasians (Table 1 and data table S6). Mitochondrial Haplogroup K has so far not been found in Paleolithic hunter-gatherers^*19*^. However, Y-haplogroup C1a2 has been reported in some of the earliest European hunter-gatherers^*17,* *20,* *21*^. The early farmers belong to common early Neolithic mitochondrial (N1a, U3 and K1a) and Y chromosome types (C and G2a), with the exception of the Levantine BAJ001 which represents the earliest reported individual carrying the mitochondrial N1b group (Table 1 and data table S6).

We examined alleles related to phenotypic traits in the ancient genomes (data table S7). Notably, three of the AAF carry the derived allele for rs12193832 in the HERC2 (hect domain and RLD2) gene that is primarily responsible for lighter eye color in Europeans^*22*^. The derived allele is observed as early as 14,000 -13,000 years ago in individuals from Italy and the Caucasus^*17,* *23*^ but had not yet been reported in early farmers or hunter-gatherers from the Near East.

## Discussion

By analyzing genome-wide-data from pre- and early-Neolithic Anatolians and Levantines, we describe the demographic developments leading to the formation of the Anatolian early farmer population that later replaced most of the European hunter-gatherers and represents the largest ancestral component in present-day Europeans^*4,* *5*^.

We report a long-term persistence of the local Anatolian hunter-gatherer gene pool over seven millennia and throughout the transition from foraging to farming. This demographic pattern is similar to those previously observed in earlier farming centers of the Fertile Crescent^*6*^ and differs from the pattern of the demic diffusion-based spread of farming into Europe^*4,* *5*^. Our results provide a genetic support for archaeological evidence^*3*^ suggesting that Anatolia was not merely a stepping stone in a movement of early farmers from the Fertile Crescent into Europe but rather a place where local hunter-gatherers adopted ideas, plants and technology that led to agricultural subsistence.

Interestingly, while the local population structure remains highly stable, a pattern of genetic interactions with neighboring regions is observed from as early as the Late Pleistocene and into the early Holocene. External genetic contributions, associated with two distinct early farming populations of the Fertile Crescent, substituted about 10% and 20% of indigenous ancestry each. The earlier one is associated with Neolithic Iran/Caucasus and the later one with Neolithic Levant. Wide temporal gaps between available genomes currently limit our ability to distinguish the mode of transfer. Obtaining additional genomic data from these regions as well as from geographically intermediate populations of eastern Anatolia and the greater Mesopotamia region could help determine how these gene flows were introduced into central Anatolia: e.g., whether by a short-term massive migration or a low-level background gene flow in an “isolation by distance” manner.

To the west, a genetic link is observed between the Anatolian and Southeastern European Pleistocene hunter gatherers. Inspection of the shared genetic components between these two populations provides us with a model explaining the genetic affinity between late Pleistocene Europeans and present-day Near Eastern populations^17^. Further sampling in Anatolia and Southeastern Europe is needed to specify the spatiotemporal extent of the genetic interactions that we observe.

## Materials and Methods

### aDNA analysis

We extracted and prepared DNA for next generation sequencing in two different dedicated ancient DNA facilities (Liverpool and Jena).

#### Liverpool, UK

Sampling and extraction steps for the individuals from Pinarbaş1 and Boncuklu were carried out in the aDNA labs at Liverpool John Moores University. The outer layer of the bone was removed using powdered aluminium oxide in a sandblasting instrument. Then, the bone was UV irradiated for 10 minutes on each side and ground into fine powder using a cryogenic grinder Freezer/Mill. DNA was extracted from 100 mg of bone powder following an established protocol^*8*^. The extracts were then shipped to Jena, Germany where downstream processing was performed.

#### Jena, Germany

All pre-amplification steps were performed in dedicated aDNA facilities of the Max Planck Institute for the Science of Human History (MPI-SHH). The inner ear part of the petrous bones of the individuals from Kfar HaHoresh and Ba’ja was sampled by drilling^*24*^ and DNA was extracted from 76 - 109 mg of the bone powder. An extraction of ∼100 mg pulverized bone from the Pinarbaş1 individual ZBC was done in the Jena facility in addition to the Liverpool extraction (the sequenced data from the two extracts of individual ZBC were merged in downstream analysis after passing the quality control step). All extractions followed the same protocol as cited for Liverpool. A 20 μl aliquot from each extract was used to prepare an Illumina double stranded, double indexed DNA library following established protocols^*9,* *25*^. Deaminated cytosines that result from DNA damage were partially removed using uracil-DNA glycosylase and endonuclease VIII but still retained in terminal read positions as a measure of aDNA authentication^*26*^. Indexed libraries were amplified using Herculase II Fusion DNA polymerase following the manufacturer’s protocol and used for two previously published downstream in-solution enrichments: a protocol targeting 1,237,207 genome-wide SNPs (‘1240k capture’^*10*^) and one targeting the entire human mitochondrial genome^*27*^. Both the initial shotgun and target-enriched libraries were single-end sequenced on an Illumina Hiseq 4000 platform (1 x 75 + 8 + 8 cycles). Sequenced reads were demultiplexed allowing one mismatch in each index and further processed using EAGER (v 1.92.54)^*28*^. First, adapter sequences were clipped and reads shorter than 30 bp were discarded using AdapterRemoval (v 2.2.0)^*29*^. Adapter-clipped reads were subsequently mapped with the BWA aln/samse programs (v 0.7.12)^*30*^ against the UCSC genome browser’s human genome reference hg19 with a lenient stringency parameter (“- n 0.01”). We retained reads with Phred-scaled mapping quality scores ≥ 20 and ≥ 30 for the whole genome and the mitochondrial genome, respectively. Duplicate reads were subsequently removed using DeDup v0.12.2^*28*^. Pseudo-diploid genotypes were generated for each individual using pileupCaller which randomly draws a high quality base (Phred-scaled base quality score ≥ 30) mapping to each targeted SNP position (https://github.com/stschiff/sequenceTools). To prevent false SNP calls due to retained DNA damage, two terminal positions in each read were clipped prior to genotyping. The genotyping produced between 129,406 to 917,473 covered targeted SNPs and a mean coverage ranging between 0.16 and 2.9 fold per individual (Table 1).

#### Dataset

We merged the newly reported ancient data and data reported by *Mathieson et al. 2017* ^*18*^ with a dataset that has been described elsewhere^*6*^. This dataset includes 587 published ancient genomes^*6,* *7,* *10,* *11,* *14,* *17,* *20,* *23,* *31-34*^ and genomes from 2,706 individuals, representing world-wide present-day populations^*6,* *35*^ that were genotyped on the Affymetrix Axiom^TM^ Genome-Wide Human Origins 1 array^*4*^ (‘HO dataset’) with a total of 597,573 SNP sites in the merged dataset. To minimize bias from differences in analysis pipelines, we re-processed the raw read data deposited for previously published Neolithic Anatolian genomes^*7*^ (labeled Tepecik_pub and Boncuklu_pub) in the same way as described for the newly reported individuals.

#### aDNA authentication and quality control

We estimated authenticity of the ancient data using multiple measures. First, blank controls were included and analyzed for extractions as well as library preparations (Data table S8). Second, we assessed levels of DNA damage in the mapped reads using mapDamage (v 2.0)^*36*^. Third, we estimated human DNA contamination on the mitochondrial DNA using schmutzi^*37*^. Last, we estimated nuclear contamination in males with ANGSD (v 0.910)^*38*^, which utilizes haploid X chromosome markers in males by comparing mismatch rates of polymorphic sites and adjacent ones (that are likely to be monomorphic). The genetic sex of the reported individuals was determined by comparing the genomic coverage of X and Y chromosomes normalized by the autosomal average coverage. To avoid bias caused by grouping closely related individuals into a population, we calculated the pairwise mismatch rates of the Boncuklu individuals following a previously reported method^*39*^ (Data table S9).

Five of the twelve individuals reported here were excluded from the population genetic analysis: two due to a high genomic contamination level (> 5 %) and three due to low amount of analyzable data (< 10,000 SNPs covered); (Data table S1).

#### Principal component analysis (PCA)

We used the smartpca software from the EIGENSOFT package (v 6.0.1)^*40*^ with the lsqproject option to construct the principal components of 67 present-day west Eurasian groups and project the ancient individuals on the first two components (fig. S4).

#### ADMIXTURE analysis

We used ADMIXTURE (v 1.3.0)^*41*^ to perform an unsupervised clustering of 3293 ancient and present-day individuals in the HO merged dataset, allowing the number of clusters (k) to range between 2 and 20. Pruning for linkage disequilibrium (LD) was done by randomly removing one SNP from each pair with genotype r^2^ ≥ 0.2, using PLINK (v 1.90)^*42,* *43*^; (--indep-pairwise 200 25 0.2). The analysis was replicated five times for each k value with random seeds and the highest likelihood replicate is reported (fig. S1 and S5) Five-fold cross-validation errors were calculated for each run.

#### D-statistics

To estimate allele frequency correlations between populations, *D*-statistics were computed using the *qpDstat* program (v 701) of the ADMIXTOOLS package^*44*^ (v 4.1) with default parameters. In order to determine whether a test population is symmetrically related to populations X and Y, the *D*-statistic *D* (*X, Y; Test, Outgroup*) was used. In particular, when comparing the affinity of different European hunter-gatherers to Near-Eastern ones in the *D*-statistic of the form *D* (*European HG1, European HG2; Near Eastern HG, Outgroup*), both the central African *Mbuti* and the Altai Neanderthal (*Altai_published.DG*) were used to check if the differing level of Neanderthal ancestry in these hunter-gatherers affects the results. Otherwise, Mbuti was used as the single outgroup. The above statistics are reported when more than 30,000 SNP positions were overlapping between the four analyzed populations.

#### Modeling ancestry proportions

We used the qpWave (v400) and qpAdm (v 632) programs of ADMIXTOOLS^*6,* *12*^ to test and model admixture proportions in a studied population from potential source populations (reference populations). As the explicit phylogeny is unknown, a diverse set of outgroup populations (Supplementary Information sections 1.2-1.4) was used to distinguish the ancestry of the reference populations.

For estimating admixture proportions in the tested populations, we used a basic set of seven outgroups including present-day populations (Han, Onge, Mbuti, Mala, Mixe) that represent a global genetic variation and published ancient populations such as Natufian^*6*^, that represents a Levantine gene pool outside of modern genetic variation and the European Upper Palaeolithic individual Kostenki14^*20*^. As a prerequisite for the admixture modeling of the target population we tested whether the corresponding set of reference populations can be distinguished by the chosen outgroups using qpWave^*6*^ (Supplementary text S3). In some cases, when a reference population did not significantly contribute to the target in the attempted admixture models, it was removed from the reference set and added to the basic outgroup set in order to increase the power to distinguish the references. In cases where “Natufian” was used as a reference population, we instead used the present-day Near-Eastern population “BedouinB” as an outgroup.

For estimations of Basal Eurasian ancestry, we followed a previously described qpAdm approach^*6*^ that does not require a proper proxy for the Basal Eurasian ancestry, which is currently not available in unadmixed form. This framework relies on the basal phylogenetic position of both Basal Eurasian and an African reference (the ancient Ethiopian *Mota* genome^*45*^) relative to other non-Africans. Thus, by using a set of outgroups that includes eastern non-African populations (Han; Onge; Papuan) and Upper Palaeolithic Eurasian genomes (*Ust’-Ishim*^*46*^, *Kostenki14, MA-1*^*47*^) but neither west Eurasians with detectable basal Eurasian ancestry nor Africans, the mixture proportion computed for *Mota* (α) can be used indirectly to estimate the Basal Eurasian mixture proportion of west Eurasian populations.

#### Mitochondrial DNA analysis

The endogenous mitochondrial consensus sequences were inferred from the output of schmutzi^*37*^, using its log2fasta program and a quality cutoff of 10. Mitochondrial haplotypes were established by aligning these consensuses to rCRS^*48*^ using the online tool haplosearch^*49*^. The coverage of each of the reported SNPs was confirmed by visually inspecting the bam pileup in Geneious (v11.0.4)^*50*^. The resulting consensus sequences were then analyzed with HaploFind^*51*^ and Haplogrep^*52*^ to assign mitochondrial haplogroups and double-checked with the rCRS oriented version of Phylotree^*53*^.

#### Y-chromosome analysis

To assign Y-chromosome haplogroups we used yHaplo^*54*^. Each male individual was genotyped at 13,508 ISOGG consortium SNP positions (strand-ambiguous SNPs were excluded) by randomly drawing a single base mapping to the SNP position, using the same quality filters as for the HO dataset. In addition to the yHaplo automated haplogroup designations, we manually verified the presence of derived alleles supporting the haplogroup assignment.

#### Phenotypic traits analyses

We tested for the presence of alleles related to biological traits that could be of interest in the geographical and temporal context of the reported ancient populations, including lactase persistence^*55,* *56*^, Malaria resistance^*57,* *58*^, Glucose-6-phosphate dehydrogenase (G6PD) deficiency^*59,* *60*^ and skin pigmentation^*23,* *61,* *62*^. The allele distribution for the SNP positions listed in Data table S7 was tabulated for each individual using Samtools mpileup (v 1.3).

#### Carbon dating

The phalanx bone from individual ZBC (Pinarbaş1) and the petrous bone from individual KFH2 (Kfar HaHoresh) were each sampled and directly radiocarbon dated at the CEZ Archaeometry gGmbH, Mannheim, Germany (table S1). Collagen was extracted from the bone samples, purified by ultrafiltration (fraction >30kD), freeze-dried and combusted to CO_2_ in an Elemental Analyzer (EA). CO_2_ was converted catalytically to graphite. The dating was performed using the MICADAS-AMS of the Klaus-Tschira-Archäometrie-Zentrum. The resulting ^14^C ages were normalized to d13C=-25% ^*63*^ and calibrated using the dataset INTCAL13^*64*^ and the software SwissCal 1.0^*65*^.

## Data availability

Alignment data (BAM) is deposited in the European Nucleotide Archive (ENA) under the accession numbers (Study PRJEB24794).

## Acknowledgments

We thank G. Brandt, A. Wissgott, F. Aron, M. Burri, C. Freund, and R. Stahl (MPI-SHH) for their support in laboratory work, M. Oreilly for graphic support, A. Gibson for help in proofreading and the members of the population genetics group in the Department of Archaeogenetics, SHH-MPI for their input and support

### Funding

This work was supported by the Max Planck Society. Experimental work at LJMU and LR stay at the MPI-SHH were funded with an internal grant from LJMU Faculty of Science to EFD. The funding for the Ba‘ja project was granted by the German Research Foundation (GZ: 80 1599/14-1) and ex oriente e.V., Berlin. Kfar HaHoresh fieldwork was supported (to NGM) by the Israel Science Foundation (Grants 840/01, 558/04, 755/07, 1161/10), The National Geographic Society (Grant 8625/09) and The Irene Levi Sala CARE Foundation. The Konya Plain fieldwork was funded by The British Institute in Ankara, British Academy (Research Development Award BR100077), a British Academy Large Research Grant LRG 35439, Australian Research Council (grants DP0663385 and DP120100969), National Geographic award GEFNE 1-11, University of Oxford (Wainwright Fund), Australian Institute for Nuclear Science and Engineering (AINSE awards AINGRA05051 and AINGRA10069), Wenner-Gren Foundation for Anthropological Research (Postdoctoral Research Grant 2008 The Origins Of Farming In The Konya Plain, Central Anatolia), Institute for Field Research

### Author contributions

J.K., E.F.D., I.H., C.J.,W.H. and M.F. conceived the study. D.B., A.F., G.M., J.P., I.H., H.M., N.G.M., M.B. and J.G. provided archaeological material. D.B., J.P., A.F., G.M., I.H., H.M., N.G.M., M.B., J.G. and P.W.S advised on the archaeological background and interpretation. D.B., J.P., E.F.D., N.G.M., M.B. and J.G. wrote the archaeological and sample background sections. M.F., L.R. and R.A.B. preformed the laboratory work. M.F., E.F.D., L.R., C.P. and C.J. preformed data analysis. M.F., E.F.D., C.J. and J.K. wrote the manuscript with input from all coauthors

### Competing interests

The authors declare no competing interests

### Data and materials availability

The genomic alignment data (BAM format) is available through the European Nucleotide Archive (ENA) under the accession numbers (Study PRJEB24794).

